# Five Decades of Rariphotic Research: A Systematic Review of Trends, Biases, and Future Directions

**DOI:** 10.64898/2026.07.07.736978

**Authors:** Almog Haim, Gal Eyal

**Affiliations:** The Goodman Faculty of Life Sciences, Bar Ilan University, Ramat Gan, Israel; The Interuniversity Institute for Marine Sciences, Eilat, Israel; School of the Environment, The University of Queensland, St. Lucia, Australia

**Keywords:** Rariphotic zone, Deep-reef ecosystems, Marine technology, Biodiversity exploratory, Systematic review

## Abstract

The rariphotic zone, typically spanning depths of approximately 130 to 300 meters, represents a key transition between light-dependent coral reef ecosystems and the aphotic deep sea. Despite its potential ecological importance, including its proposed role as a refuge for species exposed to climate-driven stress, rariphotic ecosystems remain poorly understood. In this study, we conducted a systematic review and synthesis of the scientific literature on these habitats from 1970 to 2025. Following the PRISMA 2020 protocol, we analyzed 185 studies to characterize the historical development of research, identify geographic and methodological biases, and assess shifts in research priorities over five decades.

Our results show a marked increase in research effort over the last decade, driven in part by advances in underwater technologies such as Remotely Operated Vehicles (ROVs), Human Occupied Vehicles (HOVs), and Baited Remote Underwater Video Station (BRUVS). However, this growth remains uneven, with persistent biases toward benthic rather than pelagic studies and a strong concentration of research in geographically accessible regions. Multivariate analyses of research novelty indicate that technological innovation and the formal recognition of the rariphotic zone in 2018 corresponded with major structural shifts in literature. Although the rariphotic zone is now increasingly recognized as an ecologically distinct component of the reef continuum, it remains underrepresented in ecological theory and conservation frameworks. Future research should move beyond descriptive taxonomic mapping toward integrative, data-driven functional ecology, with particular emphasis on long-term monitoring and depth-stratified connectivity.

## Introduction

Coral reefs worldwide are facing unprecedented ecological crises, driven primarily by climate change, ocean warming, and anthropogenic activities (Bellwood et al., 2004; Hoegh-Guldberg, 2011; Hughes et al., 2017, 2003; Liu et al., 2012). While these impacts have been extensively documented in shallow-water reefs (Donner et al., 2005; Hasan, 2018; Hoegh-Guldberg et al., 2007; Mai, 2024; Richmond et al., 2018), our understanding of the ecosystems located further down the seascape slope remains limited. Over the last two decades, propelled by the “deep reef refugia” hypothesis (Glynn, 1996) and advancements in diving and imaging technologies, there has been a significant surge in research on Mesophotic Coral Ecosystems (Hinderstein et al., 2010; Kahng and Kelley, 2007; Laverick et al., 2018; Loya et al., 2016). Spanning roughly from 30 to over 150 meters in depth, these areas have proven to be complex, highly diverse habitats that may offer a refuge for species stressed in shallower waters.

Beyond the lower limits of the mesophotic zone lies a cryptic depth band that remains largely unexplored: the rariphotic zone (*rarus* = scarce). Generally defined as the depth range where light penetration is insufficient to support photosynthesis-driven reef building (approximately 130 to 300 meters) (Baldwin et al., 2018), this zone represents a unique transition zone. Beyond light availability, the rariphotic zone is geomorphologically defined by the transition between the continental shelf and the continental slope. A key feature of this zone is the shelf break, which typically occurs at approximately 200 m depth (Emery, 1969). This major boundary, along with complex geomorphological structures such as submarine canyons and seamount peaks, acts as a critical conduit for organic matter and influences local hydrodynamics, potentially creating biological hotspots in an otherwise nutrient-limited environment (Brun et al., 2023; Fernandez-Arcaya et al., 2017; Genin et al., 1986; Huang et al., 2014; Palomino et al., 2011; Rogers, 2008). It serves as a transitional boundary between the light-dependent upper reef and the aphotic deep-sea environment (Jacquemont et al., 2024; Stefanoudis et al., 2019b, 2019a). The rariphotic zone is characterized by more extreme environmental conditions, lower temperatures, higher hydrostatic pressure, and minimal light availability (Baldwin et al., 2018), yet it supports remarkably diverse biological communities, including novel fish assemblages, sponges, ahermatypic corals (Cedeño-Posso et al., 2022; Reed, 2004; Rogers, 1999), and structural benthic invertebrates (Stefanoudis et al., 2019b).

Despite growing recognition of the rariphotic zone^’^s ecological significance, our current knowledge remains primarily descriptive, fragmented, and often biased. Unlike easily accessible shallow reefs, deep-reef exploration logistical and technological accessibility are more challenging. Historically, our understanding was limited by the operational depths of SCUBA, but the integration of Remotely Operated Vehicles (ROVs), Baited Remote Underwater Video Systems (BRUVS), Human-Occupied Vehicles (HOVs), and molecular tools such as environmental DNA (eDNA) (Robertson et al., 2022; Sih et al., 2019; Tolimieri et al., 2020), has begun to unveil the cryptic diversity of these depths. However, each method carries inherent biases from extractive sampling to non-invasive imaging which significantly influence the ecological data collected and our subsequent interpretation of community structure. Consequently, collected data are typically derived from rapid, localized snapshot surveys rather than continuous monitoring (Cui et al., 2023; Harvey et al., 2007).

Currently, the bulk of research effort is still concentrated in the “exploration” phase, characterized by the basic identification and mapping of new species (Baldwin et al., 2016b; Chimienti et al., 2025; Sparks et al., 2021; Tea et al., 2025). Although the continuous description of novel taxa underscores the uniqueness and underexplored nature of these deep-reef systems, the transition from basic taxonomic mapping to a comprehensive understanding of ecological function and system dynamics is significantly delayed. There is a glaring gap in our knowledge regarding community interactions, trophic structures, and how biological assemblages respond to environmental drivers across large depth gradients. To move beyond descriptive exploration, it is essential to address the systemic research biases that currently leave entire ecological components undocumented. Future efforts must shift towards integrative, functional studies that evaluate the resilience, connectivity, and complex dynamics of organisms inhabiting the rariphotic zone.

To address these knowledge gaps, this systematic review provides a quantitative and qualitative synthesis of the scientific literature concerning deep-reef habitats and rariphotic zones over five decades (1970 – 2025). Specifically, the aims of this review are to:

1. Map the scope and geographic distribution of research efforts, identifying underrepresented regions and ecosystems.
2. Critically evaluate systemic biases in biological focus, comparing research effort dedicated to pelagic versus benthic communities, and examining the influence of habitat type (e.g., hard vs. soft substrate).
3. Analyzing how the evolution of methodological tools (from SCUBA to ROVs, HOVs, BRUVS and molecular techniques) has shaped the ecological questions that can be investigated.
4. Outline essential directions for future research, emphasizing the necessary shift from discovery-based studies to data-driven functional ecology.

## Method

### Search Strategy and Information Sources

To provide a comprehensive synthesis of rariphotic zone research, a systematic literature review was conducted in accordance with the PRISMA 2020 (Preferred Reporting Items for Systematic Reviews and Meta-Analyses) guidelines (Page et al., 2021). The primary bibliographic search was performed across two major scientific databases, Web of Science and Scopus, covering publications from 1970 to 2025. However, a significant challenge in this field is that the term “rariphotic” is relatively new, having been formalized only recently (Baldwin et al., 2018). Consequently, standardized keyword indexing in major databases often fails to capture the full historical breadth of research conducted within this depth range. The search strings utilized Boolean operators (MacFarlane et al., 2022) to link geographic/depth keywords with methodological tools and biological targets. For Web of Science, the query utilized Topic Tags (TS), while the Scopus search targeted Titles, Abstracts, and Keywords. The core search terms included: (rariphotic OR “deep reef*” OR “lower mesophotic” OR “deep coral reef*”) AND (“ROV” OR “remotely operated vehicle” OR “AUV” OR “autonomous underwater vehicle” OR “HOV” OR “human occupied vehicle” OR “BRUVS” OR “baited remote underwater video systems”, “eDNA”, or “environmental DNA”) AND (“fish” OR “fish community” OR “benthic” OR “benthic community” OR “sessile” OR “sessile community”). To overcome this limitation, the initial database queries were supplemented by an ^’^active search^’^ using a snowballing technique. This involved a manual screening of the reference lists of all identified seminal papers to retrieve historical legacy studies, such as early submersible surveys, that predated current rariphotic nomenclature. This dual approach ensured that the review remains comprehensive across temporal scales: standardized database queries effectively captured modern technological and molecular-based studies (e.g., eDNA), while the manual snowballing successfully identified foundational historical records.

### Screening and Eligibility Criteria

The screening process followed a rigorous three-phase protocol. First, all retrieved records were aggregated, and duplicates were removed using bibliographic management software. Second, titles and abstracts were screened to ensure direct relevance to rariphotic ecosystems. Studies focused exclusively on shallow-water reefs (<30 m) or strictly aphotic deep-sea environments were excluded. The final phase involved a comprehensive full-text review to confirm eligibility based on depth and biological reporting. While the rariphotic zone is typically defined as 130–300 meters (Baldwin et al., 2018), this review applied an expanded depth criterion of 85 to 350 meters. This decision was made to capture studies focusing on the “lower-mesophotic” and “upper-rariphotic” ecotones, ensuring a continuous understanding of community shifts across these transitional boundaries.

### Data Extraction and Categorization

For each included study, data was extracted and recorded in a standardized database. Temporal dataincluded the publication year, further categorized into decadal bins to facilitate trend analysis. Geographic coverage was classified by bioregion, while the ecological context focused on habitat substrate (Hard, Soft, or Open Water) and the targeted community (Pelagic, Benthic, or both).

Methodological frameworks were categorized by the primary research tool employed (e.g., ROV, HOV, BRUVS, Trawl net, Acoustic records, or Water sampling for molecular analysis). Furthermore, research methods were classified by their primary approach, ranging from biodiversity surveys to environmental characterization. Finally, each study was assigned to primary research motivation, such as Biodiversity Exploratory, Ecological Function, or Conservation, to evaluate the shifting priorities of the scientific community over time.

### Data Analysis and Temporal Categorization

The majority of the data in this review are presented through descriptive synthesis, utilizing frequency distributions and proportional breakdowns to visualize research effort across regions, tools, and habitats, these visualizations were generated in R (v. 4.5.0). To facilitate the identification of long-term trends and mitigate the impact of annual fluctuations in publication volume, the data were aggregated into decadal (e.g., 1980–1989, 1990–1999). The final temporal category (2020–2025) was defined as a six-year period to capture the most recent advancements in the field. This binning approach allows for a clearer comparison of how methodological preferences and research motivations have shifted across distinct eras of deep-reef exploration.

To move beyond descriptive counts and evaluate the qualitative evolution of the field, a Multivariate Novelty Index (Pandolfi et al., 2020) was calculated. This index utilizes a Bray-Curtis dissimilarity matrix to compare the “research profile” (the specific combination of tools, regions, methods, and motivations) of each year against all preceding years. Years with an index approaching 1.0 represent “Novel Research Communities”, points in time where the scientific framework shifted significantly due to the introduction of new technologies or the exploration of unprecedented ecological questions. Other than this multivariate index, no complex statistical models were applied, as the primary goal of the visualizations is to provide a transparent representation of the current state of knowledge.

## Results

A total of 185 studies were included in the final synthesis (Fig. 1). The literature on the rariphotic zone increased over the last five decades, with relatively few studies in the earliest decades and a marked rise in publication output in the most recent years (Table 1).

**Table 1.** Decadal distribution of publications within the rariphotic zone knowledge base (1970–2025), showing individual and cumulative contributions to the total number of included studies (n = 185).

| Decadal | Number of publications (n) | Percentage (%) | Cumulative Percentage (%) |
| --- | --- | --- | --- |
| 1970-1979 | 5 | 2.7% | 2.7% |
| 1980-1989 | 22 | 11.9% | 14.6% |
| 1990-1999 | 25 | 13.5% | 28.1% |
| 2000-2009 | 36 | 19.5% | 47.6% |
| 2010-2019 | 51 | 27.6% | 75.2% |
| 2020-2025 | 46 | 24.9% | 100% |

**Figure 1.**
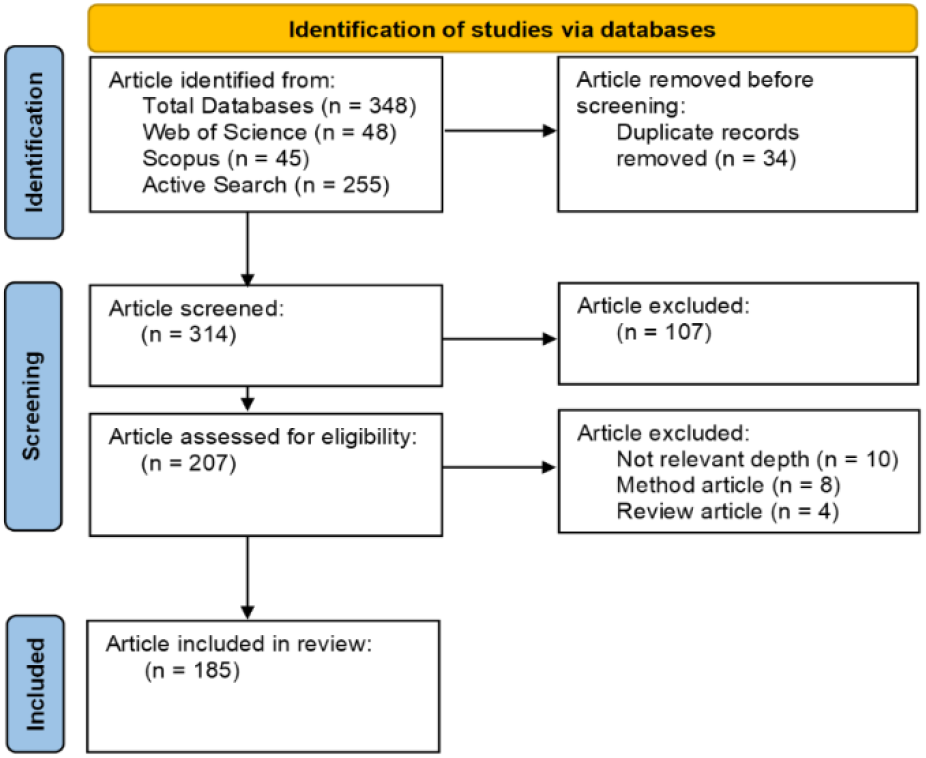
PRISMA 2020 Flow Diagram - A flow diagram illustrating the systematic literature review process according to the PRISMA 2020 standards. The figure details the identification, screening, and inclusion of studies (n = 185)related to rariphotic ecosystems.

### Temporal trends

The temporal distribution of studies shows a steady increase in research effort over time (Table 1). Only 5 studies were identified in 1970 – 1979 (2.7% of the total dataset), followed by 22 studies in 1980 – 1989 (11.9%), 25 studies in 1990 – 1999 (13.5%), 36 studies in 2000 – 2009 (19.5%), 51 studies in 2010 – 2019 (27.6%), and 46 studies in 2020 – 2025 (24.9%). The highest absolute number of publications occurred during 2010 – 2019, while the most recent period already accounts for nearly one quarter of the total dataset despite covering only six years.

### Geographic distribution and motivations

Research effort was unevenly distributed across bioregions, with some regions receiving substantially more attention than others (Fig. 2A). Research motivations also varied across the dataset, with discovery-oriented studies being the most common category, followed by studies focused on ecological function and conservation (Fig. 2B).

**Figure 2.**
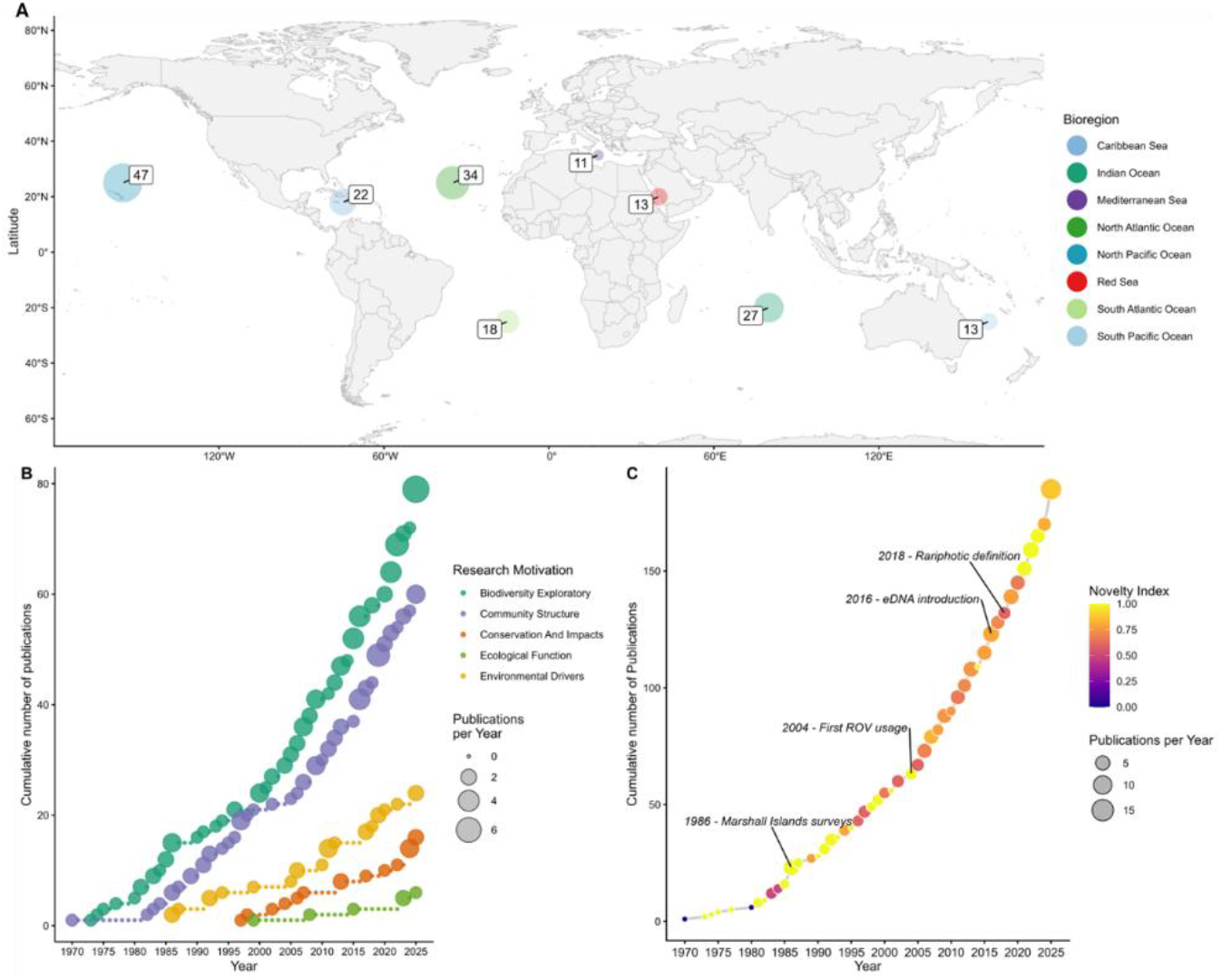
Spatiotemporal Dynamics and Research Novelty - **(A)** Global distribution of rariphotic research sites across bioregions, with bubble sizes proportional to the number of publications. **(B)** Proportional representation of research motivations (e.g., Biodiversity Exploratory, Ecological Function) across five decades. **(C)** Cumulative growth of rariphotic literature, color-coded by the Novelty Index, showcasing the emergence of ‘novel research communities’ following technological shifts and the formal definition of the rariphotic zone (2018).

### Research novelty

The multivariate novelty analysis quantified temporal shifts in the structural composition of rariphotic literature. The Novelty Index identified several distinct periods where the research profile (combinations of tools, regions, methods, and motivations) significantly diverged from preceding years, with values reaching or approaching 1.0 (Fig. 2C). Notable peaks in novelty were recorded in 1986, 2004, 2016 and 2019, marking points of significant framework emergence.

### Methodological structure

The methodological composition of the literature was dominated by a limited set of survey and sampling tools, including ROVs, BRUVS, HOVs, trawls, acoustic approaches, and water sampling for molecular analyses (Fig. 3A). Research methods were dominated by survey-based studies, followed by sample collection,environmental characterization, and molecular analysis (Fig. 3B).

**Figure 3.**
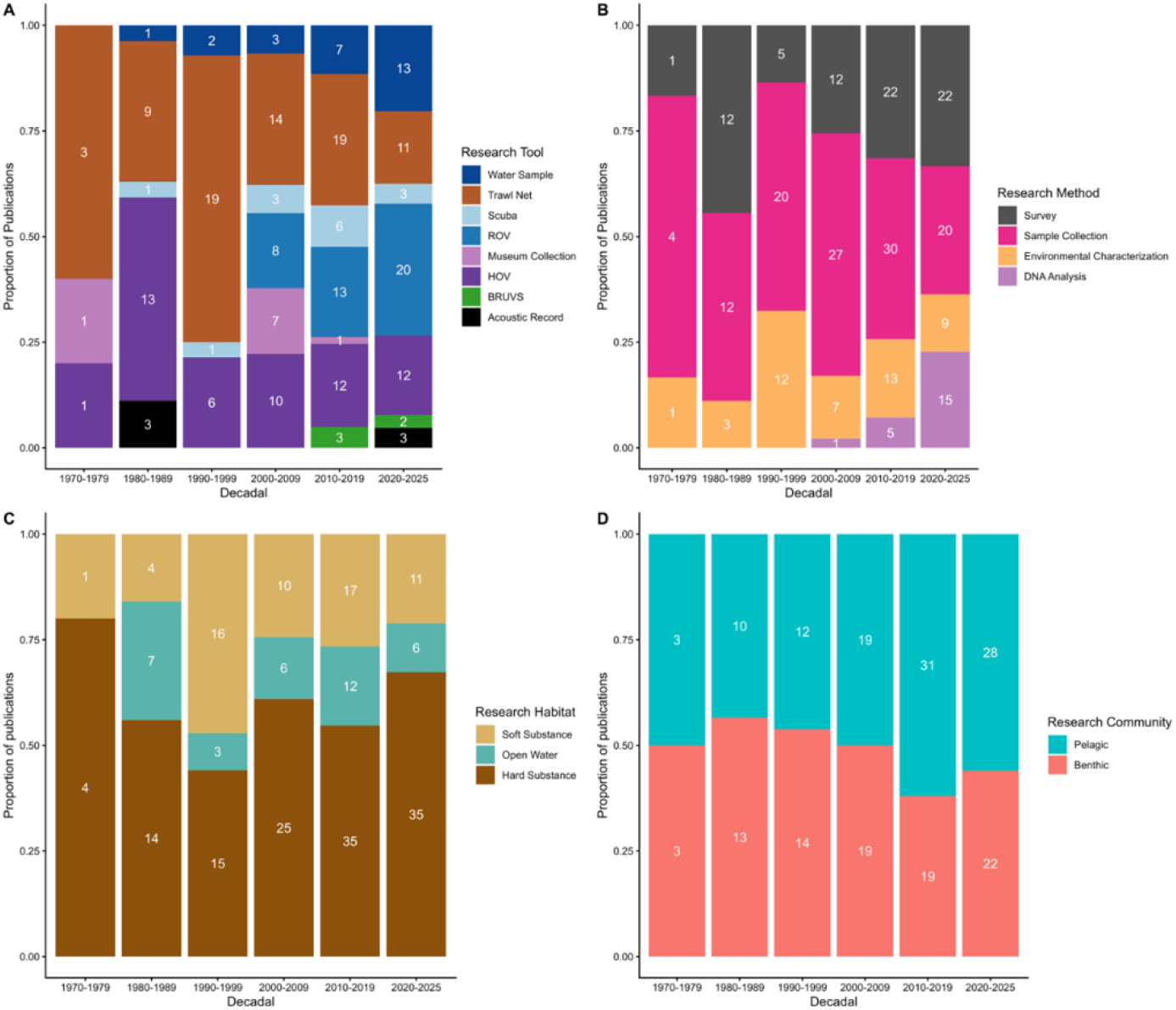
Methodological and Environmental Landscape – A multivariate characterization of the rariphotic knowledge base. The panels provide a breakdown of: **(A)** Technical tool (e.g., ROVs, BRUVS, HOVs), **(B)** Technical method (e.g., Survey, Sample collection, eDNA), **(C)** Research habitat (e.g., hard vs. soft substrates), and **(D)** Target community focus (Benthic vs. Pelagic). ROV = Remotely Operated Vehicle, HOV = human operated Vehicle, BRUVS = Baited Remote Underwater Video Surveys.

### Habitat and community focus

Target communities were unevenly represented across the literature, with benthic-focused studies more frequent than pelagic-focused studies (Fig. 3C). Habitat representation also varied, with studies distributed unevenly among hard-bottom, soft-bottom, and open-water settings (Fig. 3D).

### Community-specific patterns

Community-specific patterns differed between benthic and pelagic studies across tools, habitats, methods, and research motivations (Fig. 4A–D). Benthic studies were more strongly associated with hard-bottom habitats and survey-based approaches, whereas pelagic studies were less frequent and more limited in methodological range.

**Figure 4.**
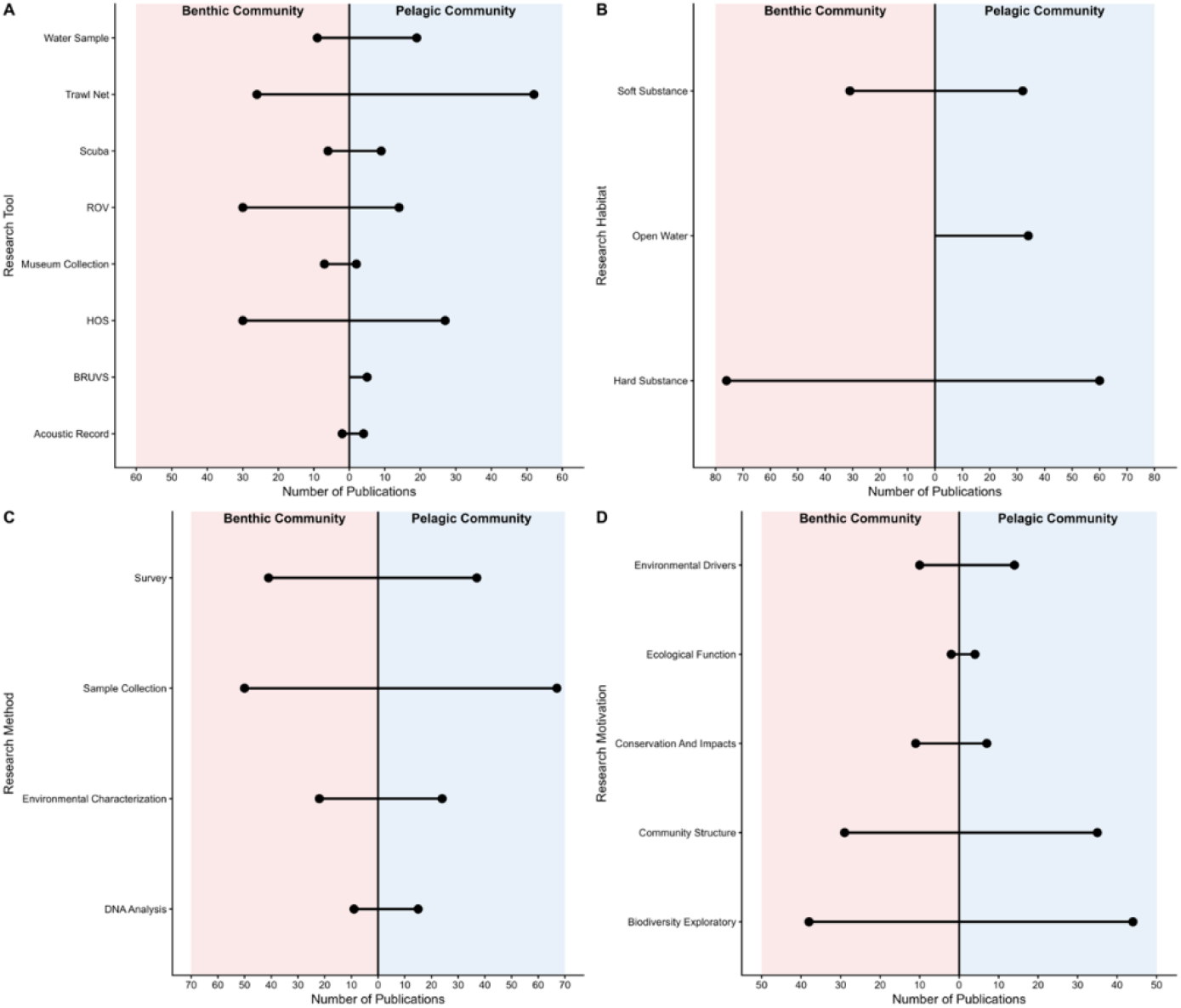
Community-Specific Research Trends **-** A comparative analysis of research effort between Benthic and Pelagic communities. The figure contrasts publication counts across different technical tools **(A)**, habitat **(B)**, method **(C)** and motivations **(D)**.

## Discussion

This review shows that rariphotic research has expanded markedly over the past five decades, yet it remains structurally uneven across regions, methods, and ecological focus. The rise in publication output is not simply a matter of more studies being produced over time; it also reflects a deeper shift in how the scientific community has begun to view deeper reef zones, moving them from the margins of reef science toward a more recognized ecological domain. At the same time, literature still carries clear signs of its exploratory origins, with a strong concentration of work in a relatively small number of accessible regions and a heavy reliance on descriptive survey-based approaches. These patterns suggest that the rariphotic zone is now widely acknowledged as important, but it is still far from being understood with the same depth or resolution as shallower reef systems. Beyond quantitative growth, the multivariate novelty analysis reveals that the evolution of rariphotic research has been characterized by discrete qualitative ^’^pulses^’^ that restructured the field’s conceptual framework. These shifts align with the concept of ^’^Novel Research Communities^’^ (Pandolfi et al., 2020), where the integration of new technologies and geographic targets creates unprecedented ecological frameworks. For instance, the high novelty observed in 1986 corresponds to the geographic expansion of deep-reef surveys into underrepresented Pacific bioregions, such as the Marshall Islands (Colin et al., 1986; Maragos and Jokiel, 1986; Thresher and Colin, 1986). Subsequently, the 2004 novelty peak marks a pivotal technological shift with the initial integration of Remotely Operated Vehicles (ROVs) into rariphotic exploration (Grigg, 2004; Quattrini et al., 2004; Reed, 2004). More recently, the introduction of molecular tools in 2016, specifically environmental DNA (eDNA) metabarcoding, added a new methodological dimension to biodiversity assessments (Baldwin et al., 2016a, 2016b; Thomsen et al., 2016). The surge in 2019 followed the formal definition of the rariphotic zone in 2018 (Hsieh and Lo, 2019; Khalaf et al., 2019; Stefanoudis et al., 2019b, 2019a; Tamir et al., 2019; Weijerman et al., 2019). Our results indicate that the field is not merely growing in volume but is undergoing a state of flux, characterized by rapid discovery pulses rather than a standardized methodology.

A central feature emerging from this review is that progress in the field has been driven more by access and opportunity than by a fully integrated research framework. For many years, the rarity of studies was largely a consequence of technical limitations, depth constraints, and the difficulty of working in habitats beyond conventional SCUBA depth ranges (Danovaro et al., 2014; Roger, 2015; Smith et al., 2022). As new platforms, imaging systems (Bell et al., 2022), and molecular tools became available (Duhamet et al., 2023; Yamahara et al., 2025), research activity increased and the scope of questions broadened (Glover and Smith, 2003). However, the literature still shows that certain components of rariphotic ecosystems, particularly pelagic assemblages, functional processes, and less accessible geographic regions, remain underrepresented. This imbalance matters because it limits the extent to which the rariphotic zone can be interpreted as a coherent ecological system rather than as a patchwork of locally sampled habitats.

The overall picture is therefore one of rapid but uneven development. The field has clearly advanced beyond the stage of simply documenting the existence of rariphotic communities, but it has not yet reached a stage where its biodiversity, connectivity, and ecological functioning are equally well resolved across space and taxa. In that sense, the rariphotic zone now occupies an important middle ground in reef science: it is no longer invisible, yet it is still not fully integrated into broader ecological theory or management frameworks (Baldwin et al., 2018; Stefanoudis et al., 2019b).

### Zonation and distinctiveness

The rariphotic zone should be treated as a distinct ecological transition rather than a simple extension of mesophotic reef habitats (Stefanoudis et al., 2019b). Its definition was first formalized in 2018 by Baldwin et al, based on depth distributions and the distinct fish fauna observed below the mesophotic range, establishing the rariphotic as a named reef zone rather than an arbitrary depth interval (Baldwin et al., 2018). Subsequent studies have reinforced this view by showing strong depth-related turnover in both fish and benthic communities, together with limited connectivity to shallower reefs (Jacquemont et al., 2024; Stefanoudis et al., 2019a). In particular, community composition often shifts sharply around the mesophotic– rariphotic boundary (Koslow et al., 2000; Weijerman et al., 2019), and the lower rariphotic itself appears to represent a recognizable faunal in some regions (Reed, 2004; Rogers, 1999). These findings support the idea that the rariphotic zone is ecologically discrete, with assemblages shaped by depth-associated gradients in light, habitat structure, and connectivity.

### Geographic imbalance

The geographic distribution of rariphotic research remains strongly uneven, with most studies concentrated on a relatively small number of accessible reef systems. This pattern is not surprising given the technical challenges of working at these depths, but it does mean that the current literature is biased toward regions where deep-reef exploration has been logistically feasible and historically prioritized (Fricke, 1996; Fricke and Hottinger, 1983; Fricke et al., 1987; Kaiser et al., 1993). The rariphotic zone was originally defined from work in the southern Caribbean, particularly Curaçao (Baldwin et al., 2018; Harasewych and Tëmkin, 2015; Liddell et al., 1997; Pratt et al., 2020), where repeated submersible surveys revealed a distinct deep-reef fauna and motivated the formal recognition of the zone (Bongaerts et al., 2015; Jacquemont et al., 2024). However, later studies have shown that depth zonation and community turnover can vary among regions (Chimienti et al., 2025; Stefanoudis et al., 2019b, 2019a; Swanborn et al., 2022), and that the extent of connectivity to shallower reefs is not necessarily uniform across locations. As a result, the present knowledge base likely captures only part of the global variability in rariphotic ecosystems, and expanding work into underrepresented bioregions will be essential for testing how general these patterns really are.

### Methodological bias

The methodological structure of rariphotic research has been shaped by the practical difficulty of sampling deep reef environments. Most studies in literature relied on survey-based (Bongaerts et al., 2015; Busby et al., 2005; Colin, 1974; Fricke and Meischner, 1985; Hissmann et al., 1998; Mejía-Quintero et al., 2022; Stefanoudis et al., 2025; Tamir et al., 2019) or specimen-based approaches (Bullis and Struhsaker, 1970; Gj∅saeter, 1984; Khalaf and Zajonz, 2007; Merten et al., 2021; Pires, 1992; Pochon et al., 2015; Rocha et al., 2024; Rogers, 1999), which are effective for documenting biodiversity but are also sensitive to access limitations, observer effects, and habitat visibility. This has likely contributed to a strong bias toward organisms and habitats (Hughes et al., 2021) that are easier to detect with visual or opportunistic methods, particularly in benthic settings. The growing use of molecular tools such as eDNA offers a way to complement these approaches and reduce some of their taxonomic blind spots, especially for cryptic or mobile taxa that are often underrepresented in conventional surveys (Iguchi et al., 2024; Sawh et al., 2025). However, these methods also introduce their own sources of bias through sampling design, DNA recovery, amplification, and downstream filtering, which means that methodological integration rather than replacement is likely the most productive path forward (Cote et al., 2023; van der Loos and Nijland, 2021).

### Benthic versus Pelagic

The imbalance between benthic and pelagic studies reflects a broader focus on substrate-associated communities in rariphotic research. Benthic assemblages have been shown to vary sharply across depth, with distinct communities occupying shallow, mesophotic, and rariphotic habitats and with relatively low connectivity among these depth zones (Chimienti et al., 2025; Stefanoudis et al., 2019b). This has encouraged a strong emphasis on benthic habitat characterization and on the spatial turnover of reef-associated taxa, particularly in studies using visual surveys and direct sampling. By contrast, pelagic communities remain comparatively poorly represented, despite their importance for energy transfer, trophic structure, and benthic-pelagic coupling in reef systems. The resulting imbalance leaves an important gap in our understanding of how rariphotic ecosystems function as whole systems, rather than as substrate-bound assemblages alone.

### Conservation and refuge hypothesis

The conservation relevance of the rariphotic zone is closely tied to the deep reef refuge hypothesis (Glynn, 1996), which proposes that deeper reefs may buffer shallow reef communities from disturbance and provide sources of recolonization after impact (Bridge et al., 2013; Hughes and Tanner, 2000; Lesser et al., 2009; West and Salm, 2003). The recognition of the rariphotic as a distinct deep-reef zone initially raised interest in its possible role as a refuge for shallow-water species facing warming, degradation, and other anthropogenic pressures. However, studies examining depth-stratified reef assemblages have shown that rariphotic communities are often compositionally distinct from shallower reefs and may have limited connectivity to them, which complicates the assumption that deeper habitats can function as universal refuges (Jacquemont et al., 2024; Stefanoudis et al., 2019b, 2019a; Weijerman et al., 2019). At the same time, more recent work has shown that human impacts and fishing pressure can extend into deeper depth ranges, meaning that rariphotic ecosystems are not automatically protected from disturbance (Asher et al., 2017; Grigg, 2004; Roblet et al., 2025; Stefanoudis et al., 2025). These findings suggest that the conservation value of the rariphotic zone is substantial, but context-dependent, and should be evaluated on the basis of regional connectivity, depth-specific pressure, and community composition rather than assumed a priority.

### Future directions

Future research on the rariphotic zone should move beyond descriptive biodiversity surveys and toward integrative approaches that can address function, connectivity, and ecosystem dynamics across depth gradients. A major priority is the expansion of sampling into underrepresented regions, since the current literature is still concentrated in a relatively small number of accessible reef systems. In parallel, greater use of complementary methods, including molecular tools, environmental characterization, and standardized survey protocols, would help reduce methodological bias and improve comparability across studies. This is especially important for resolving cryptic diversity and for linking benthic and pelagic processes, both of which remain poorly understood in rariphotic ecosystems. Long-term monitoring would also be valuable, particularly in the context of warming, fishing pressure, and other disturbances that increasingly extend into deeper reef environments. More broadly, future work should aim to test whether rariphotic habitats function as refugees, transition zones, or independently structured communities, rather than assuming that they serve the same ecological role everywhere. Such a shift would help place the rariphotic zone within a more explicit functional and conservation framework.

## Acknowledgements

We are grateful to Reem Neri for his invaluable assistance in refining the systematic search strings. We also thank Mika Doron and Liat Biniuri for their support with graphical analyses.

## Notes

### Competing Interest Statement

The authors have declared no competing interest.

